# An Explainable Knowledge Graph-Driven Approach to Decipher the Link Between Brain Disorders and the Gut Microbiome

**DOI:** 10.64898/2025.12.11.693657

**Authors:** Naafey Aamer, Muhammad Nabeel Asim, Sebastian Vollmer, Andreas Dengel

**Author notes:** These authors contributed equally to this work (Naafey Aamer, Muhammad Nabeel Asim).

## Abstract

**Motivation:** The communication between the gut microbiome and the brain, known as the microbiome-gut-brain axis (MGBA), is emerging as a critical factor in neurological and psychiatric disorders. This communication involves complex pathways including neural, hormonal, and immune interactions that enable gut microbes to modulate brain function and behavior. However, the specific mechanisms through which gut microbes influence brain function remain poorly understood, and existing computational efforts to understand these mechanisms are simplistic or have limited scope.

**Results:** This work presents a comprehensive approach for elucidating the interactions that allows gut microbes to influence brain disorders. We construct a large curated biomedical knowledge graph comprising 586,318 nodes across 16 entity types and 3,573,936 edges spanning 103 relation types, integrating ontological and experimental data relevant to the MGBA. On this graph, we train GNN-GBA, a GraphSAGE-based graph neural network with a DistMult relation-aware decoder, achieving an AUC-ROC of 0.997 and an F1-score of 0.981 on link prediction, outperforming nine baseline methods across four categories. Using GNNExplainer, we extract and rank mechanistic pathways connecting gut microbes to brain disorders, and demonstrate their stability across multiple random initializations. GNN-GBA successfully identified pathways for 125 brain disorders, revealing shared metabolite hubs (including flavonoids, bile acids, and short-chain fatty acids) that mediate gut–brain communication across diverse neurological conditions. Furthermore, we show that the top pathways are consistent with existing literature for three common disorders. Lastly, we develop an interactive dashboard (GutBrainExplorer) to explore thousands of potential mechanistic pathways across 125 brain disorders, which is publicly available at https://sds-genetic-interaction-analysis.opendfki.de/gut_brain/.

**Availability:** Code and data are available at https://github.com/naafey-aamer/GNN-GBA.

## Introduction

Brain disorders impose a staggering global burden, they are the leading cause of disability and the second leading cause of death worldwide [20]. As shown in Figure 1, these conditions are highly complex and fall into diverse categories, including developmental, immune-mediated, neurodegenerative, and psychiatric disorders [58]. Some of these common and highly recurring conditions have no treatment options like Alzheimer’s disease, Parkinson’s disease, major affective disorder, epilepsy, and schizophrenia [40]. The mechanisms behind brain disorders are often multifactorial, reflecting the interplay of genetic, environmental, and, as recent studies suggest [35, 51, 45, 36], microbial factors.

**Fig. 1.**
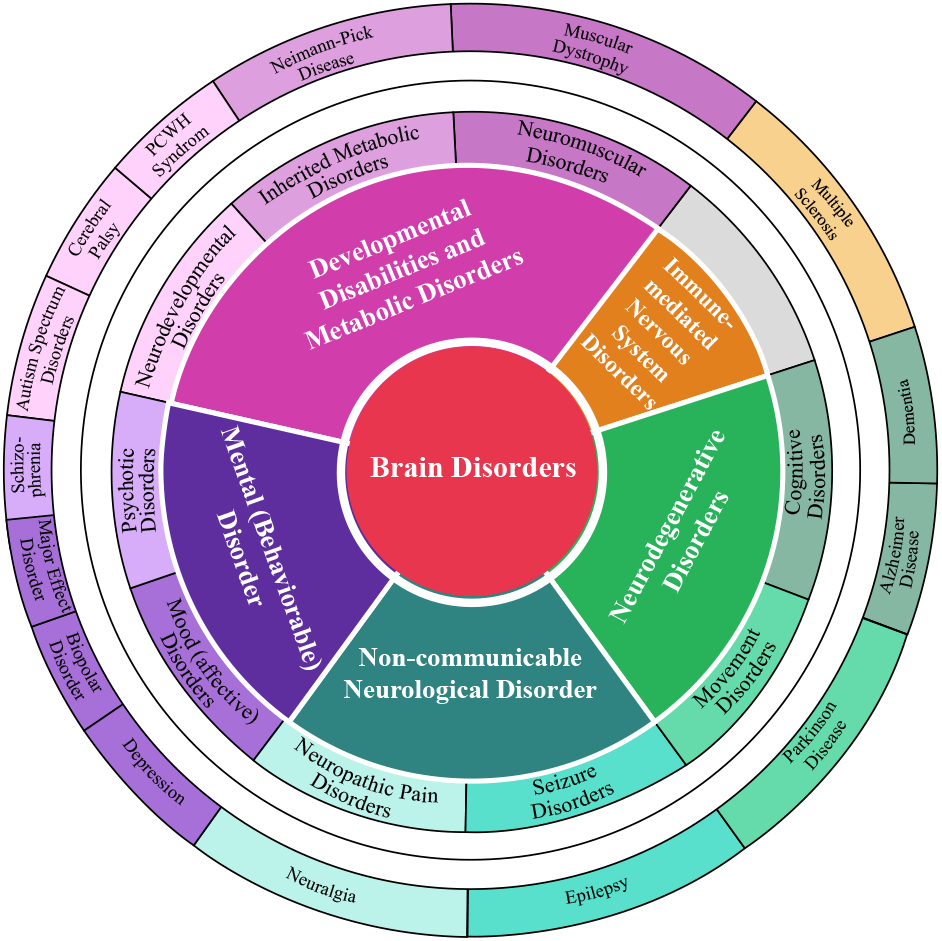
Brain disorders classified into several categories and sub-categories.

The relationship between brain disorders and these microbial factors is not fully understood [2], and it may be attributed to interactions between intestinal microbes and the central nervous system, which is known as the microbiome-gut-brain axis (MGBA). The MGBA is a communication network linking the gut and brain via neural, endocrine, immune and metabolic pathways [40] and a growing body of studies implicate it in neurological conditions [14, 40]. These findings suggest that the gut microbiome may be a key factor in neurodegeneration and neurodevelopment.

Central to gut-brain communication are gut metabolites, small bioactive chemicals generated during microbial metabolic processes of the digestive system [47]. These metabolites are critical messengers between the gut and the brain [47], binding to receptors on various cells in both systems to modulate key processes such as inflammation and neurotransmission [16, 8]. Beyond their role as messengers, these metabolites can influence metabolism and gene expression, resulting in functional mechanistic pathways between gut microbes and various diseases [63]. Without a clear understanding of these mechanistic pathways, our comprehension of gut microbes’ role in disease remains incomplete, hindering the development of targeted therapeutic strategies. Since the gut microbiome composition is influenced by diet, medications and other confounding factors[14], existing studies often struggle to take a global view of this intricate network of cause and effect, making it challenging to develop a comprehensive understanding of these interactions. Traditional analytical methods typically examine one data modality at a time and tend to miss complex, multi-scale interactions. There is therefore a critical need for an integrative approach that can jointly analyzes diverse types of microbial, genetic, metabolic and phenotypic data.

Biomedical knowledge graphs offer a powerful solution for such integrative analysis [43]. These graphs are constructed by merging diverse data types into structured, machine-readable formats [9, 68]. They are able to represent biomedical entities (genes, proteins, microbes, diseases, metabolites, etc.) as nodes and their relationships (e.g. molecular interactions, causal links, phenotypic associations) as edges [43]. To explore the MGBA, the inclusion of metabolites is particularly crucial, as they represent the functional output of microbial activity and serve as the link between microbes and brain function. Graph machine learning has been highlighted as a key emerging tool in biomedicine, with the potential to accelerate biomedical discovery [32]. Once, the biomedical graph is constructed, graph machine learning can be directly applied to it to learn the complex interaction patterns in the MGBA.

This paper explores the mechanisms between brain disorders and the gut microbiome through biological knowledge graphs and explainable graph neural networks. Our contributions include:

1. **A curated biomedical knowledge graph** focused on the MGBA, comprising 586,318 nodes (spanning microbes, metabolites, proteins, genes, diseases, and 11 additional entity types) and 3,573,936 edges covering 103 distinct biological relation types, constructed by integrating the PheKnowLator ontology pipeline with the GutMGene database.
2. **An explainable graph neural network (GNN-GBA)** that combines a 3-layer GraphSAGE encoder with a DistMult relation-aware decoder to learn the complex biological interactions within this knowledge graph. GNN-GBA achieves an AUC-ROC of 0.997 and F1-score of 0.981, substantially outperforming nine baseline methods spanning knowledge graph embeddings (TransE, DistMult, ComplEx, RotatE), relation-aware GNNs (R-GCN, CompGCN), attention-based GNNs (GAT), and shallow embedding approaches (Node2Vec with Random Forest, XGBoost, and SVM). We further report per-relation performance across all 103 relation types with macro and micro averaging to demonstrate robust prediction quality even under significant class imbalance.
3. **A systematic investigation into 125 distinct brain disorders**, identifying mechanistic pathways through which specific gut microbes influence neurological conditions via metabolite-mediated cascades. Using GNNExplainer with path-level scoring, we rank these pathways and demonstrate their stability across multiple random seeds (Jaccard overlap of top-3 paths = 0.926). Centrality analysis of merged pathway graphs reveals shared hub metabolites that serve as conserved mediators of gut–brain communication. We also show that the top pathways for 3 common disorders are consistent with existing literature.
4. **An interactive dashboard** (GutBrainExplorer) that enables researchers to explore and visualize thousands of mechanistic pathways for 125 brain diseases, displaying step-by-step connections from microbes to target diseases.

This work reveals novel insights into how specific microbes might influence neurological conditions and also provides a rigorously validated, generalizable framework for exploring complex disease–microbiome interactions. The identified shared metabolite hubs suggest that the MGBA operates through a relatively small set of conserved metabolic pathways, opening avenues for targeted dietary and therapeutic interventions. The interactive dashboard promotes further investigation in this emerging field, offering a starting point for future experimental validation and therapeutic development.

## Related Work

Earlier studies have often employed conventional machine learning on microbiome profiles, for instance, Su et al. introduce a genus-level random forest ranks microbial species by importance [56]. Such models can identify important microbial features for diseases (potential “biomarkers”), but they typically treat the microbiome as an independent high-dimensional feature vector, lacking explicit representation of mechanistic pathways. To incorporate network structure, researchers have turned to graph-based methods [39]. Since the gut-brain axis information can be efficiently represented using Knowledge Graphs (KGs), multiple works have aimed to construct them. MiKG was developed to identify potential associations between gut microbiota, neurotransmitters, and mental disorders by manually extracting relevant entities and relationships from scientific publications [34]. Additionally, established biomedical databases, such as MetaCyc [31], Reactome [13], gutMGene [11], and KEGG [30] serve as valuable resources for KG construction, as they have collected diverse data on pathways, drugs, genes, and microbes. The Pre-/Probiotics Knowledge Graph (PPKG) integrates both scientific publications and existing biomedical databases to construct a comprehensive KG to store the relationship between the central nervous system and the enteric (gut) nervous system [33]. Beyond constructing KGs that encapsulate known relationships, there is growing interest in discovering novel MGBA pathways through predictive modeling and meta-search. Several graph based models have been proposed to predict microbe–disease links. For example, MGMLink leverages the gutMGene database to identify the shortest mechanistic pathways (i.e., gut-brain links) through breadth-first search (BFS) and template-based searches in a heterogeneous graph [53]. MGMLink uses node2vec [24] and TransE [5] as embedding methods which are static, precomputed, and shallow representations that learn local patterns within the knowledge graph.

More advanced works employ graph neural networks (GNNs), which have shown the promise of interpretability by highlighting important microbial neighbors that drive a given disease association [21][37]. While existing GNN-based methods like [37] and [28] have potential, they lack interpretability, and the data incorporated is upto two orders of magnitude smaller than the one used in this work. This results in limited generalizability and an inability to capture the complexity of microbe-brain interactions across diverse biological pathways. We address this need by developing GNN-GBA, a GNN trained over a large curated gut–brain-axis knowledge graph of over 3.5 million relationships, encompassing microbes, their metabolites, host targets, and neurological diseases. By training on this rich network, the model learn global interaction patterns that link microbes to brain pathology. To make these patterns human-interpretable, this study uses a graph interpretability method, GNNExplainer, [67] which highlights the mechanistic pathways which are most critical for a predicted microbe–disease link.

## Materials and Methods

This section describes our approach to knowledge graph construction, path mining, and graph neural networks to map the complex relationships between gut microbes, metabolites, and brain diseases. Figure 4 illustrates the overall flow of the methodology.

### MGBA Knowledge Graph Construction

To create the initial MGBA knowledge graph, [52]’s methodology is employed: an ontology KG was built using the PheKnowLator [7] pipeline and combined with associations from the GutMGene database. PheKnowLator is a Python 3 library that enables construction of KGs that incorporate a wide variety of data and terminology sources, including ontologies such as the Mondo Disease Ontology (MONDO) [61], the Chemical Entities of Biological Interest Ontology (CHEBI) [19], and the Human Phenotype Ontology (HPO) [23]. The gutMGene database [11] consists of microbe-metabolite and microbe-gene assertions that occur in the host of either human or mouse. PheKnowLator constructs edges from curated ontologies while GutMGene provides experimentally derived microbe–metabolite and microbe–gene associations, conflicting assertions between the two sources are rare in practice. In cases where both sources contained edges between the same entity pair, the union of all relation types was retained. No confidence scores were assigned to edges, as the constituent databases do not uniformly provide such metadata; instead, data quality was ensured through source selection (restricting to curated databases rather than text-mined assertions) and noise reduction via ontology filtering. Ontologies unrelated to the MGBA scope (including the Plant Ontology, Food Ontology, Genotype Ontology, and Transcription Factor classifications) were removed, eliminating 1,502,361 edges and 196,148 nodes that would otherwise introduce spurious paths between microbes and brain disorders. This resulted in the final MGBA knowledge graph which consists of **586**,**318** nodes spanning 16 distinct biomedical entity types (Table 1) and **3**,**573**,**936** edges covering 103 unique predicate types. Figure 2 shows a small subgraph showcasing the connectivity between “enterococcus avium ly1”, a gut microbe, and “Autism”.

**Table 1.** Node Types and Counts in the Knowledge Graph.

| Node Type | Count | Percentage (%) |
| --- | --- | --- |
| Chemical | 150,080 | 25.60 |
| SNP | 145,156 | 24.76 |
| Protein | 96,196 | 16.41 |
| GO_Term | 43,870 | 7.48 |
| Cell_Line | 41,801 | 7.13 |
| Gene | 26,838 | 4.58 |
| Disease | 23,264 | 3.97 |
| Pathway | 16,395 | 2.80 |
| Phenotype | 16,291 | 2.78 |
| Anatomy | 14,181 | 2.42 |
| Vaccine | 6,271 | 1.07 |
| Taxonomy | 2,373 | 0.40 |
| Cell_Type | 2,368 | 0.40 |
| Microbe | 814 | 0.14 |
| Chromosome | 249 | 0.04 |
| Environmental_Conditions | 171 | 0.03 |
| **Total** | **586318** | **100.00** |

**Fig. 2.**
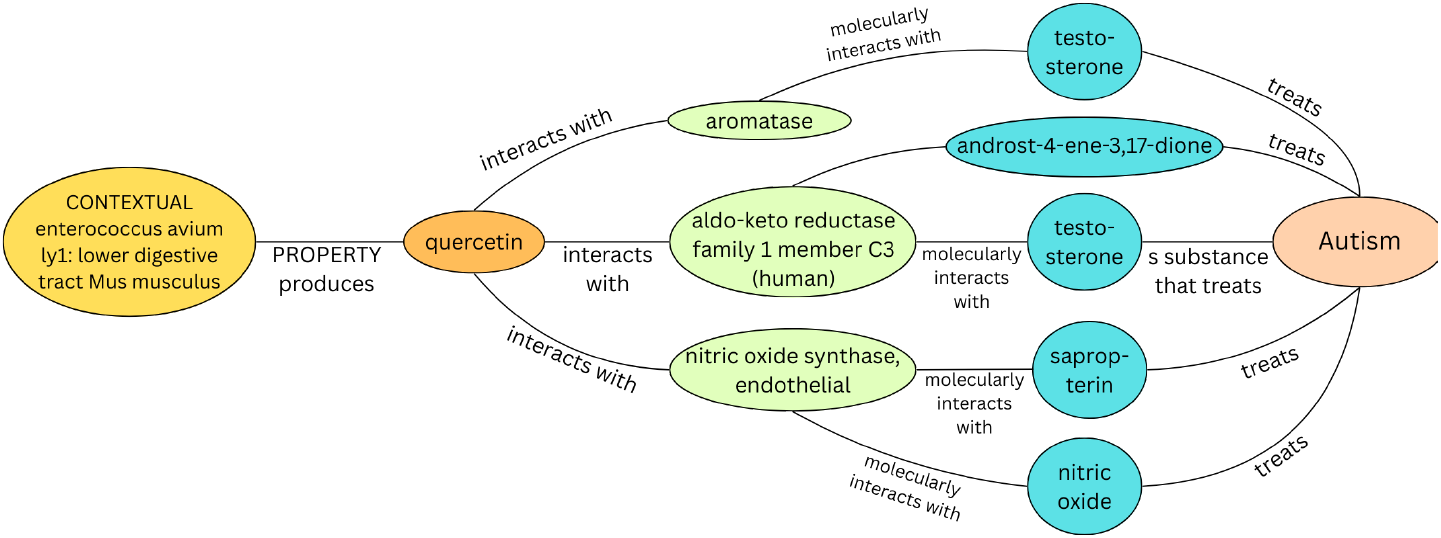
A portion of the MGBA knowledge graph with the detailed pathways from a microbe to a brain disorder.

The MGBA knowledge graph’s 103 unique predicate (relation) types have a highly skewed distribution. The five most frequent predicates are “molecularly interacts with” (961,885 edges, 26.9%), “subClassOf” (614,821 edges, 17.2%), “has phenotype” (411,610 edges, 11.5%), “participates in” (367,500 edges, 10.3%), and “is substance that treats” (268,070 edges, 7.5%). At the other end of the spectrum, 37 relation types have fewer than 1,000 edges each, including structurally specific predicates such as “conduit for” (54 edges), “proximally connected to” (54 edges), and “in right side of” (55 edges). This class imbalance across relation types motivates our use of relation-wise performance evaluation. Among these predicates, “molecularly interacts with,” “causally influences,” “causes or contributes to condition,” and “is substance that treats” are particularly valuable for tracing mechanistic pathways, as they encode direct biological causation.

It is important to note that metabolites, the critical messenger molecules between the gut and the brain, are primarily represented within the Chemical node category, which constitutes the largest node type in our graph (as shown in Table 1). This substantial representation of chemical entities, including short-chain fatty acids, neurotransmitter precursors, and bioactive peptides, enables us to trace metabolite-mediated mechanisms between gut microbes and neurological diseases.

### Disease Selection And Path Mining

To get practical insights for brain disorders, it is necessary to ensure that sufficient links exist between brain disorders and the gut microbiome in the constructed knowledge graph. For that, we initially identified 2,905 diseases associated with the brain. From this large set, the focus was on diseases with robust metabolite connections, as these are critical for understanding microbe-disease relationships. Only 138 diseases had more than 30 metabolite connections, and after grouping variants of the same disease (e.g., various Alzheimer’s disease subtypes) and removing conditions that returned zero paths with a timeout of 60 seconds, a final set of 125 neurological disorders were shortlisted for in-depth analysis. With these 125 diseases identified, paths between each disease and the 814 microbes in the knowledge graph were mined. For each disease-microbe pair, paths are searched, with the Networkx Python library [25], upto a maximum length of 5, capturing relevant connections while avoiding computational overhead. The maximum path length of 5 was selected based on both biological plausibility and empirical observation. Paths longer than 5 edges risk including indirect ontological relationships (e.g., traversing “subClassOf” hierarchies) that dilute mechanistic specificity. Empirically, the median path length across all mined disease– microbe paths was 4.0 (mean 3.93), with the vast majority falling between 3 and 4 edges, confirming that length-5 paths capture the relevant biological depth without excessive noise. Shorter cutoffs (e.g., length 3) would miss multi-step metabolite-mediated mechanisms, while longer cutoffs (e.g., length 7) did not yield additional mechanistically interpretable paths in preliminary experiments but substantially increased computational cost.

Figure 3 shows the distribution of relation types across all mined microbe–disease pathways. The three most frequent relations—”is substance that treats” (137,204 occurrences), “interacts with” (109,440), and “molecularly interacts with” (105,238)—together account for approximately 65% of all pathway edges. This indicates that the identified mechanistic pathways are primarily mediated by metabolite–protein interactions and therapeutic associations, consistent with the known role of gut metabolites as biochemical messengers in the MGBA [47]. The high frequency of “produces” (80,263) and “metabolizes” (52,486) further confirms that microbial metabolic activity is central to the pathways captured by our approach.

**Fig. 3.**
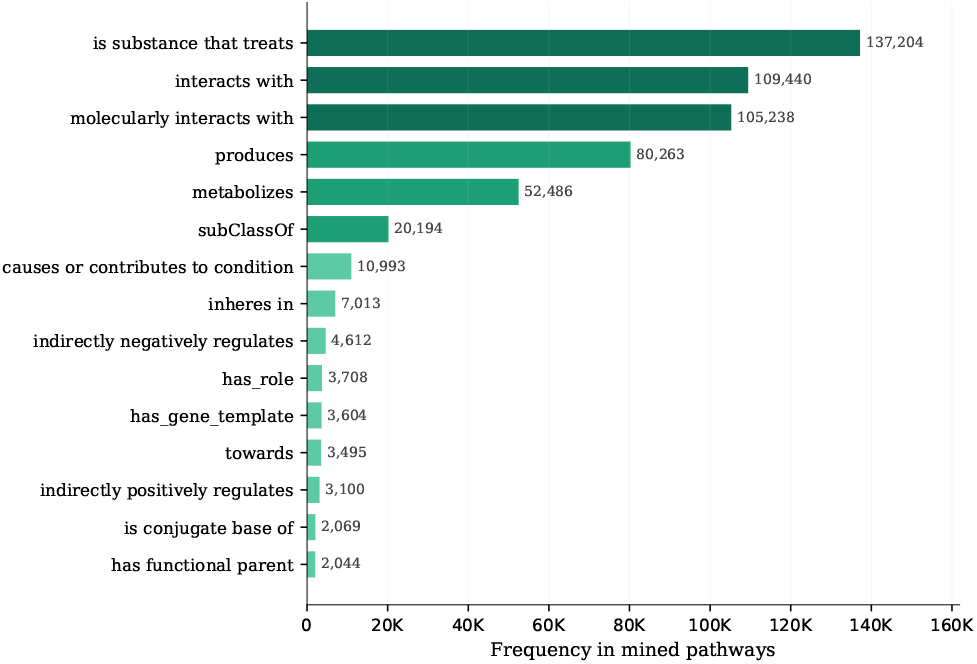
Distribution of relation types in the mined microbe–disease pathways across 125 brain disorders. The three most frequent relations— *is substance that treats, interacts with*, and *molecularly interacts with* — account for approximately 65% of all pathway edges, indicating that the identified mechanisms primarily involve metabolite-mediated molecular interactions and therapeutic associations. The prevalence of *produces* and *metabolizes* reflects the central role of microbial metabolic activity in gut– brain communication.

**Fig. 4.**
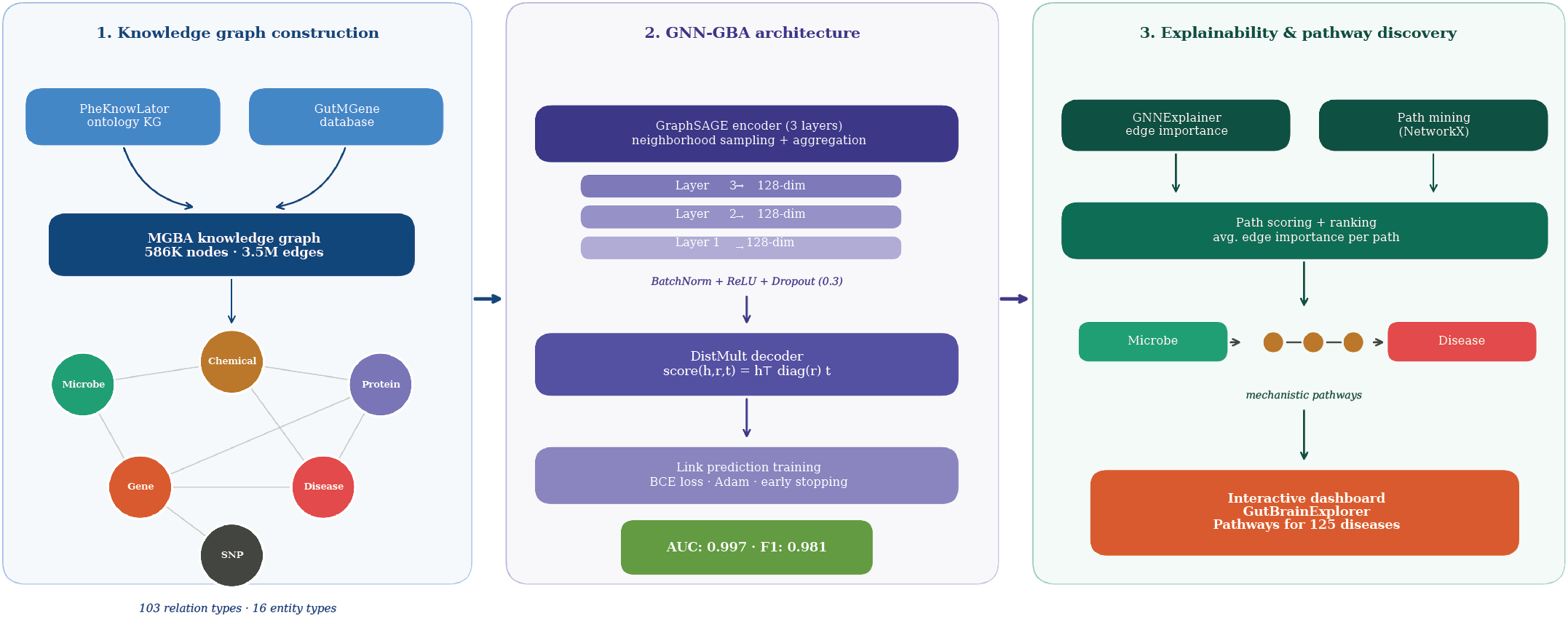
Overview of the GNN-GBA framework. **(1)** A biomedical knowledge graph is constructed by integrating the PheKnowLator ontology pipeline with the GutMGene database, resulting in 586,318 nodes across 16 entity types and 3,573,936 edges spanning 103 relation types. **(2)** GNN-GBA trains a 3-layer GraphSAGE encoder with a DistMult decoder for link prediction, achieving an AUC-ROC of 0.997 and F1-score of 0.981. **(3)** GNNExplainer and path mining are combined to identify and rank mechanistic pathways from gut microbes to brain diseases. The results are made available through the GutBrainExplorer interactive dashboard covering 125 diseases.

Analysis of the mined pathways reveals a set of metabolites that serve as shared mechanistic intermediates across a large number of brain disorders (Table 2). Notably, quercetin appears in pathways for 125 of 125 analyzed diseases, followed by arachidonic acid (124 diseases) and resveratrol (124 diseases). The dominance of flavonoids, isoflavones, and bile acids among the top intermediates suggests that the MGBA operates through a relatively small number of conserved metabolic pathways. These metabolites are well-established products of microbial metabolism [11] and have documented neuroactive properties, including anti-inflammatory, antioxidant, and neuroprotective effects [17]. The identification of these shared hubs provides both validation of our approach and actionable targets for experimental follow-up, particularly for dietary interventions that modulate flavonoid and bile acid availability in the gut.

**Table 2.**
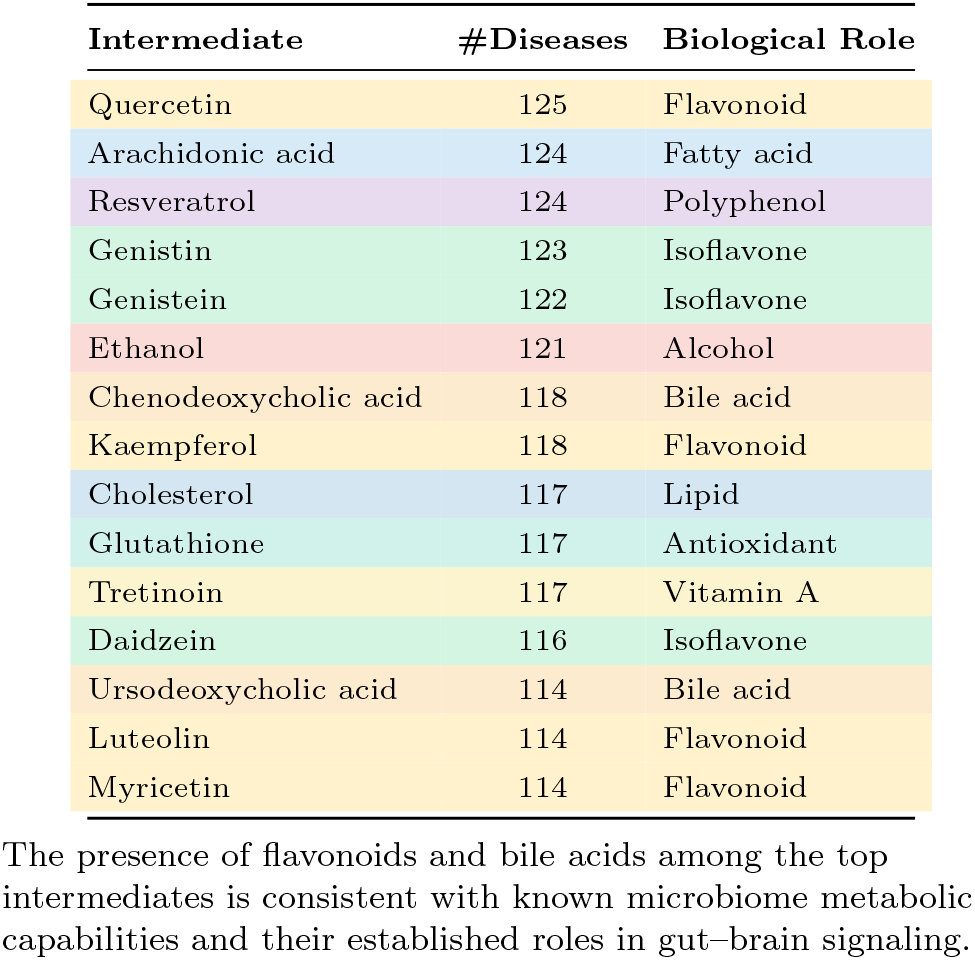
Most frequent intermediate nodes in microbe–disease pathways, representing shared mechanistic mediators through which gut microbes may influence brain disorders.

| Intermediate | #Diseases | Biological Role |
| --- | --- | --- |
| Quercetin | 125 | Flavonoid |
| Arachidonic acid | 124 | Fatty acid |
| Resveratrol | 124 | Polyphenol |
| Genistin | 123 | Isoflavone |
| Genistein | 122 | Isoflavone |
| Ethanol | 121 | Alcohol |
| Chenodeoxycholic acid | 118 | Bile acid |
| Kaempferol | 118 | Flavonoid |
| Cholesterol | 117 | Lipid |
| Glutathione | 117 | Antioxidant |
| Tretinoin | 117 | Vitamin A |
| Daidzein | 116 | Isoflavone |
| Ursodeoxycholic acid | 114 | Bile acid |
| Luteolin | 114 | Flavonoid |
| Myricetin | 114 | Flavonoid |

The presence of flavonoids and bile acids among the top intermediates is consistent with known microbiome metabolic capabilities and their established roles in gut-brain signaling.

### The GNN-GBA Architecture

To effectively model the complex relationships within the knowledge graph, GNN-GBA was trained for link prediction. It was implemented in PyTorch Geometric [22], a library that extends PyTorch [46] to handle graph-structured data efficiently. Unlike traditional methods [48, 24], GNNs simultaneously capture both the local structure (direct connections) and global context (multi-hop relationships) of the biomedical entities in the graph. This is particularly crucial for understanding the gut–brain axis, where the relationships between microbes, metabolites, and brain disorders often involve complex, multi-step pathways rather than direct links.

The GNN-GBA encoder consists of 3 GraphSAGE [26] layers, which learn node representations through iterative neighborhood sampling and aggregation. At each layer *l*, the representation of node *v* is updated as:

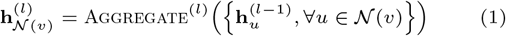

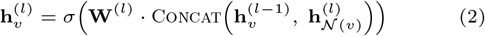

where 𝒩 (*v*) denotes the sampled neighborhood of node *v*, **W**^(*l*)^ is a learnable weight matrix, and *σ* is a nonlinear activation function. We use mean aggregation, which computes 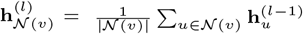.

Batch normalization is applied after each layer (except the final layer) to stabilize training, followed by ReLU activation and dropout (*p*=0.3). The encoder transforms one-hot encoded node type features into 128-dimensional embeddings that preserve the structural and semantic relationships in the knowledge graph.

We chose GraphSAGE over relation-aware GNN encoders such as R-GCN [54] or CompGCN [60] for two reasons. First, GraphSAGE’s neighborhood sampling strategy makes it scalable to our 586K-node graph, whereas R-GCN’s per-relation weight matrices become memory-prohibitive with 103 relation types at this scale. Second, we delegate relation modeling to the decoder (DistMult), which learns a separate embedding per relation type. This decoupled design allows the encoder to learn general structural patterns while the decoder specializes in distinguishing between relationship types.

For the decoding phase, GNN-GBA employs DistMult [66], a relation-aware scoring function that explicitly models the 103 different types of biological relationships in the graph. Given a candidate triple (*h, r, t*)—representing a head entity, relation, and tail entity—DistMult computes a plausibility score through bilinear scoring:

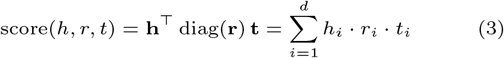

where **h, t** ∈ ℝ^*d*^ are entity embeddings produced by the GraphSAGE encoder and **r** ∈ ℝ^*d*^ is a learned relation-specific embedding (*d*=128). This element-wise trilinear product allows each dimension of the relation embedding to independently scale the interaction between corresponding dimensions of the head and tail entities, enabling the model to differentiate between mechanistic predicates (e.g., “molecularly interacts with”) and structural ones (e.g., “subClassOf”). We choose DistMult over scoring methods because of its generalizability and consistently high performance in previous work on biological knowledge graphs [1].

The model was trained using binary cross-entropy loss on both positive edges (existing relationships from the knowledge graph) and negative edges. Negative edges were generated through random corruption with a 1:1 negative-to-positive ratio: for each positive triple (*h, r, t*), either the head or tail entity was randomly replaced (with equal probability) by a uniformly sampled node to create a negative triple. To prevent training on false negatives, corrupted triples that correspond to existing edges in the knowledge graph were filtered and re-sampled. The optimization was performed using the Adam optimizer with an initial learning rate of 0.001 and weight decay of 5 *×* 10^−4^. A learning rate scheduler halved the learning rate after 5 epochs without improvement in validation AUC. Training continued with early stopping based on validation AUC, with patience of 15 epochs.

On top of the trained GNN-GBA model, a GNNExplainer [67] module was incorporated to provide explanation capabilities. GNNExplainer identifies the most important subgraph structure influencing a specific prediction by learning a soft edge mask **M** ∈ [0, 1]^|ℰ|^ that maximizes the mutual information between the original prediction and the masked prediction:

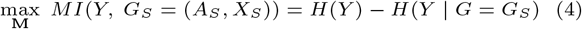

where *G*_*S*_ is the subgraph induced by the mask, *Y* is the model’s prediction, and *H*(·) denotes entropy. The mask is optimized via gradient descent for 100 epochs with a learning rate of 0.005. For each disease–microbe path, the GNNExplainer computed importance scores for individual edges within the path. These edge-level importance scores were then aggregated using average pooling to produce a single path-level score:

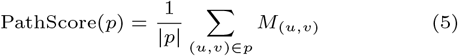

where *p* is a path (ordered sequence of edges) and *M*_(*u,v*)_ is the learned importance of edge (*u, v*). This enables systematic ranking of different pathways connecting the same microbe– disease pair, with higher-scoring paths representing more influential mechanistic routes.

### Evaluation Metrics

Model performance on the link prediction task was evaluated using five complementary classification metrics: **AUC-ROC, AUC-PRC, Precision, Recall**, and the **F1-score**. To account for the highly skewed predicate distribution in the knowledge graph, all metrics are reported both as *micro averages* (weighted by relation frequency, reflecting overall performance dominated by frequent relations) and *macro averages* (unweighted mean across all 103 relation types, giving equal weight to rare relations and revealing performance on underrepresented predicates). Macro averages are accompanied by standard deviations and 95% confidence intervals computed via the *t*-distribution across relation types.

To quantify the importance of shared mechanistic nodes from the output of the GNNExplainer, we compute two graph centrality measures using NetworkX [25]. **Betweenness centrality** measures the fraction of all shortest paths in the graph that pass through a given node, identifying nodes that serve as critical bridges between different parts of the pathway network. **PageRank** [44] computes a node’s importance based on the importance of the nodes linking to it, providing a global measure of influence that accounts for both direct connections and the broader network topology (*α*=0.85).

To assess the stability of GNNExplainer-derived pathway rankings, we run the explainer multiple times (5 seeds) for hundred disease–microbe pair and compute two complementary stability metrics. **Jaccard overlap (top-***k*) measures the fraction of shared paths among the top-*k* ranked pathways across different seeds; a Jaccard of 1.0 indicates that the same *k* paths are ranked highest in every run. **Top-***k* **consistency** reports the percentage of disease–microbe pairs for which the exact same *k* paths rank highest across all seeds, providing the most stringent measure of ranking stability. Together, these metrics distinguish between instability in the top-ranked pathways (which would undermine the reported mechanistic findings) and instability in the lower-ranked pathways (which is expected and inconsequential for the biological conclusions).

## Experimental Setup

### Data Splitting

The knowledge graph was partitioned at the edge level into training (90%), validation (5%), and test (5%) sets, resulting in 2,859,148 training triples, 357,394 validation triples, and 357,394 test triples. We used a stratified split was performed randomly at the triple level using a fixed random seed for reproducibility. Since entities may participate in multiple triples, shared entities exist across splits; this transductive setting is standard for knowledge graph link prediction [5, 66] and reflects the realistic scenario where the model predicts missing relationships between known entities. The validation set was used for early stopping and learning rate scheduling. Negative samples for evaluation were generated with a 1:1 ratio by randomly corrupting the head or tail of each positive triple, with corrupted triples filtered against the full knowledge graph to avoid false negatives.

### Baseline Models

To rigorously evaluate GNN-GBA, we compare against nine baseline methods spanning four categories, ensuring coverage of both classical and state-of-the-art approaches for knowledge graph link prediction.

Four relation-aware embedding methods were evaluated, all using 128-dimensional embeddings trained for up to 100 epochs with early stopping on validation AUC:

- **TransE** [5]: models relations as translations in embedding space (**h** + **r** ≈ **t**).
- **DistMult** [66]: uses bilinear diagonal scoring, identical to the decoder used in GNN-GBA but without a GNN encoder.
- **ComplEx** [59]: extends DistMult to complex-valued embeddings, enabling asymmetric relation modeling via Hermitian dot products.
- **RotatE** [57]: models relations as rotations in complex space, capturing symmetry, antisymmetry, inversion, and composition patterns.

These models learn entity and relation embeddings jointly but do not perform message passing, making them unable to leverage multi-hop neighborhood structure.

Two graph neural network baselines that explicitly model relation types during message passing were evaluated:

- **R-GCN + DistMult** [54]: uses relation-specific weight matrices in a 2-layer GNN encoder, paired with a DistMult decoder. This is the most architecturally comparable baseline to GNN-GBA, differing only in the encoder.
- **CompGCN + DistMult** [60]: jointly embeds nodes and relations through composition operations (subtraction) during message passing, with a DistMult decoder. Uses 128-dimensional embeddings and 2 layers.

To compare with a GNN without relation modeling, we include:

- **GAT + DistMult** [62]: uses a 2-layer Graph Attention Network encoder (4 attention heads) that learns attention-weighted aggregation without explicit relation modeling, paired with a DistMult decoder.

Three methods combine static graph embeddings with classical machine learning classifiers:

- **Node2Vec** [24] **+ Random Forest / XGBoost / SVM**: Node2Vec generated 128-dimensional embeddings using random walks of length 20, with 10 walks per node and context window of 10, trained for 10 epochs. Edge features were constructed via Hadamard products of the source and target node embeddings and fed to Random Forest [6] (*n*=100 trees), XGBoost [10] (histogram-based), and SVM [12] classifiers.

These baselines do not model relation types and treat link prediction as binary classification over edge features, providing a lower bound on performance achievable without graph neural networks or relation-aware modeling. All baseline models use 128-dimensional embeddings to ensure a fair comparison with GNN-GBA.

## Results

### Link Prediction Performance

GNN-GBA demonstrated strong performance in predicting links interactions within the MGBA knowledge graph. As shown in Table 3, GNN-GBA achieved an AUC-ROC of 0.997, AUC-PRC of 0.996, and F1-score of 0.981 on 357,394 test triples, outperforming all nine baseline methods across every evaluation metric.

**Table 3.**
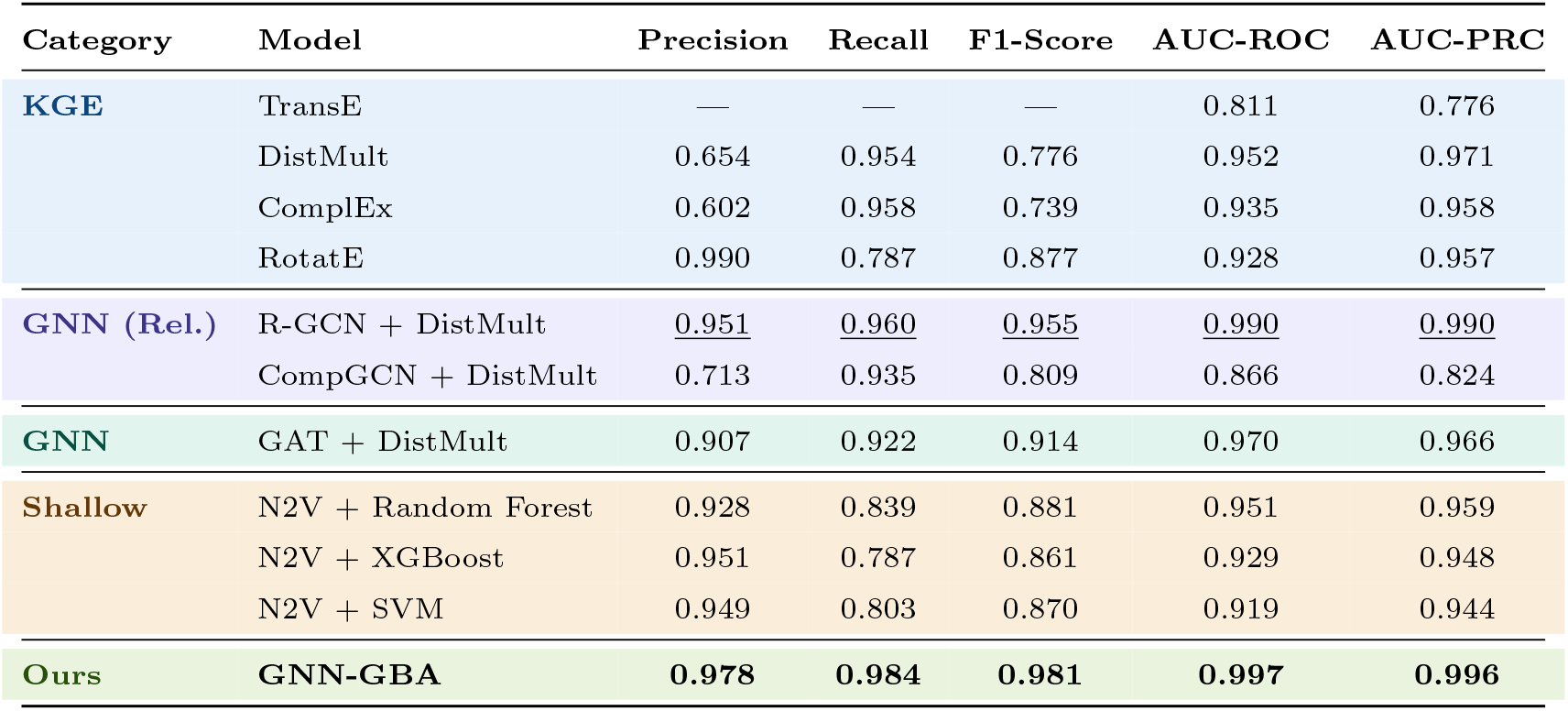
Link prediction performance comparison on the MGBA knowledge graph. **Bold** indicates the best result per metric; underline indicates second best. All models use 128-dimensional embeddings and are evaluated on 357,394 test triples.

Among the KGE baselines, DistMult achieved the highest AUC-PRC (0.971) but suffered from low precision (0.654), indicating a tendency to over-predict positive links. RotatE showed the opposite pattern, high precision (0.990) but lower recall (0.787), suggesting conservative predictions. TransE achieved an AUC-ROC of only 0.811, the lowest among all methods, and its distance-based scoring function produced values that did not cross the 0.5 decision threshold, resulting in zero precision and recall at that cutoff. ComplEx performed comparably to DistMult (AUC-ROC 0.935) but did not benefit from its asymmetric relation modeling on this graph.

The strongest baseline was R-GCN + DistMult (AUC-ROC 0.990, F1 0.955), differing from GNN-GBA only in its use use of a complex relation aware encoder. GAT + DistMult achieved an AUC-ROC of 0.970 and F1 of 0.914 without any explicit relation modeling in the encoder, demonstrating that attention-based neighborhood aggregation captures meaningful structural patterns. However, its gap relative to GNN-GBA (0.970 vs. 0.997) confirms the value of GraphSAGE’s sampling strategy for this particular graph topology.

The shallow baselines (Node2Vec + classifiers) achieved AUC-ROC values between 0.919 and 0.951. Their consistently high precision (*>*0.92) but lower recall (*<*0.84) show conservative classification behavior, while their inability to perform message passing limits their capacity to capture the multi-step pathways central to gut–brain communication.

### Per-Relation Performance Analysis

To assess whether aggregate metrics mask poor performance on specific relation types, we also evaluated GNN-GBA across all 103 predicate types individually. Figure 5(a) shows the relationship between relation frequency and performance: AUC-ROC remains consistently high (*>*0.85) regardless of relation count, while F1-score degrades sharply for rare relations with fewer than approximately 100 test edges. This frequency dependence explains the gap between the overall AUC-ROC (0.997) and the relation-wise macro-averaged F1-score (0.702).

**Fig. 5.**
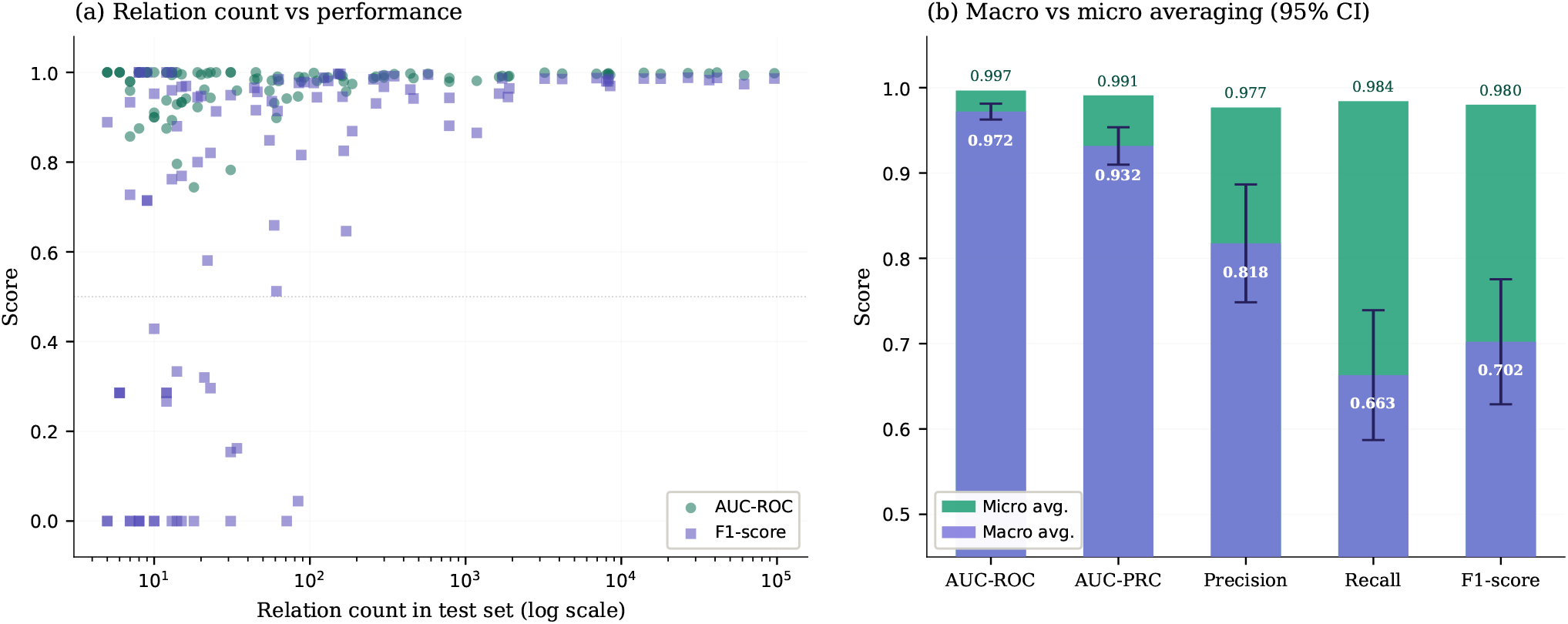
Per-relation performance analysis of GNN-GBA across 103 relation types. **(a)** Relation count (log scale) versus AUC-ROC and F1-score. Each point represents one relation type. AUC-ROC remains consistently high regardless of frequency, while F1-score degrades for rare relations with fewer than ∼100 test edges. **(b)** Overlapping macro (purple, foreground) and micro (teal, background) averages with 95% confidence intervals on macro. Where the teal bar extends above the purple, micro exceeds macro—indicating that frequent relations perform better and dominate micro averaging. The gap is largest for recall and F1-score, confirming that rare relation types are the primary source of lower macro performance. AUC-ROC shows minimal gap (0.972 macro vs 0.997 micro), confirming robust ranking ability across all relation frequencies.

Figure 5(b) quantifies this effect through overlapping macro and micro averages with 95% confidence intervals. The micro-averaged AUC-ROC (0.997) and macro-averaged AUC-ROC (0.972 *±* 0.048) are close, confirming robust ranking ability across relation types. In contrast, the gap between micro F1 (0.980) and macro F1 (0.702 *±* 0.375) is substantial, driven by 15 rare relation types (each with fewer than 10 test edges) where the model achieves an F1 of zero despite high AUC. This pattern is consistent with the known sensitivity of threshold-dependent metrics to class imbalance and does not indicate a failure of the model’s learned representations, as confirmed by the uniformly high AUC scores. The macro-averaged precision (0.818 *±* 0.353) and recall (0.663 *±* 0.389) further indicate that the primary difficulty lies in recall for rare relations, the model correctly identifies positive instances when it predicts them, but misses many true positives when training examples are scarce.

### Hub Structure Analysis

When the top-ranked pathways for each disease were analyzed, shared hub structures emerged that connect multiple diseases to various gut microbes through common mechanistic nodes (Figure 6). Figure 6 shows six representative diseases where distinct hub structures emerge, from the densely connected Landau-Kleffner syndrome to the simpler Bell’s palsy topology. To quantify these hubs, we computed betweenness centrality and PageRank for each node in the merged pathway graphs across all analyzed diseases.

**Fig. 6.**
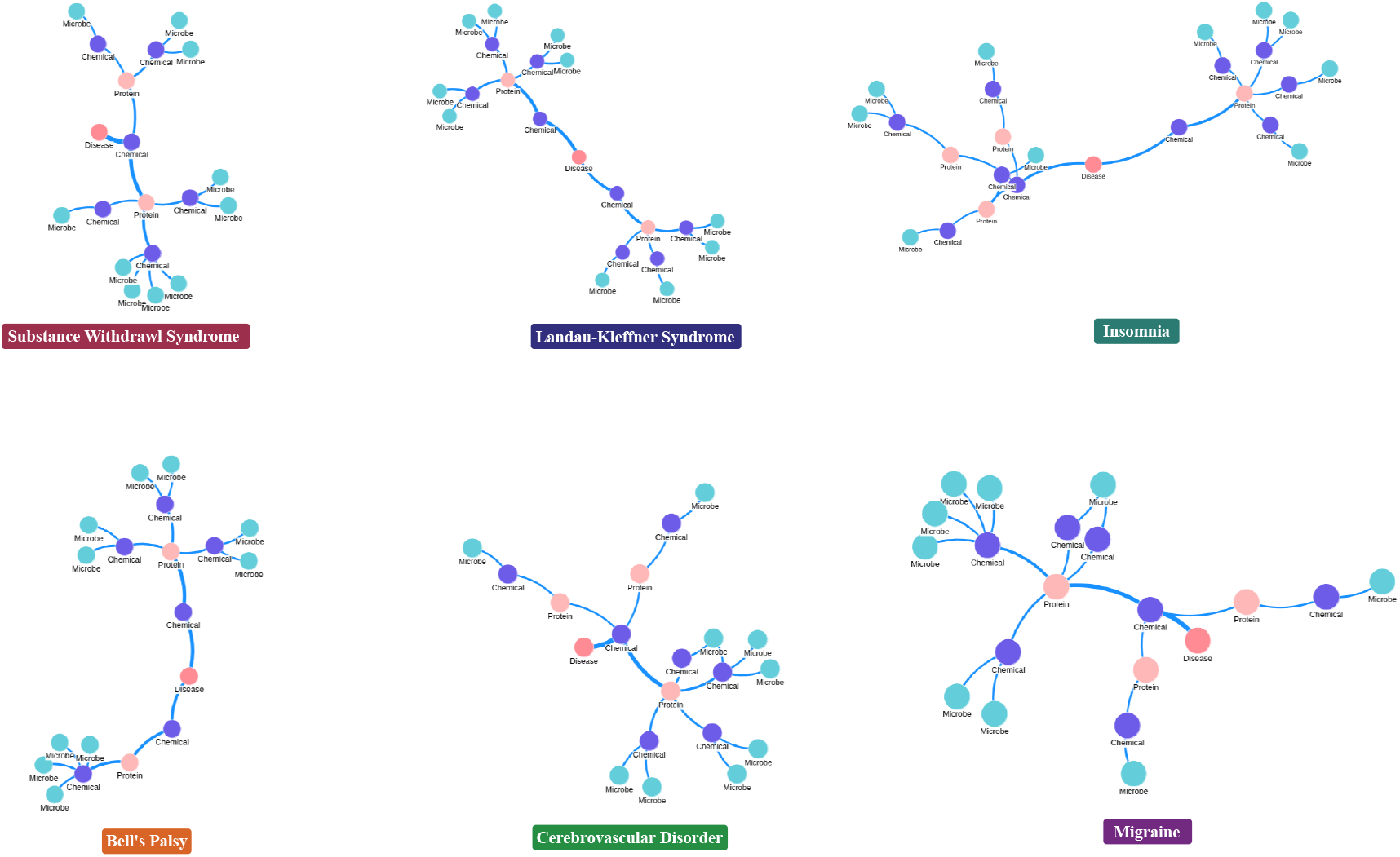
Merged top paths reveal hub structures. These hubs show common pathways through which the microbiome influences neurological health.

Table 4 presents the most prominent shared hub nodes. Levodopa bridges the most diseases (38), consistent with its established role as a dopamine precursor used across movement and psychiatric disorders [41]. Ethanol appears as a hub in 23 diseases, reflecting its well-documented neurotoxic and neuromodulatory effects. Among the metabolite hubs, cholesterol (12 diseases, avg. PageRank 0.184, avg. betweenness 0.039) and lactic acid (14 diseases) are both established products of microbial metabolism with known roles in neuroinflammation and blood–brain barrier integrity.

**Table 4.**
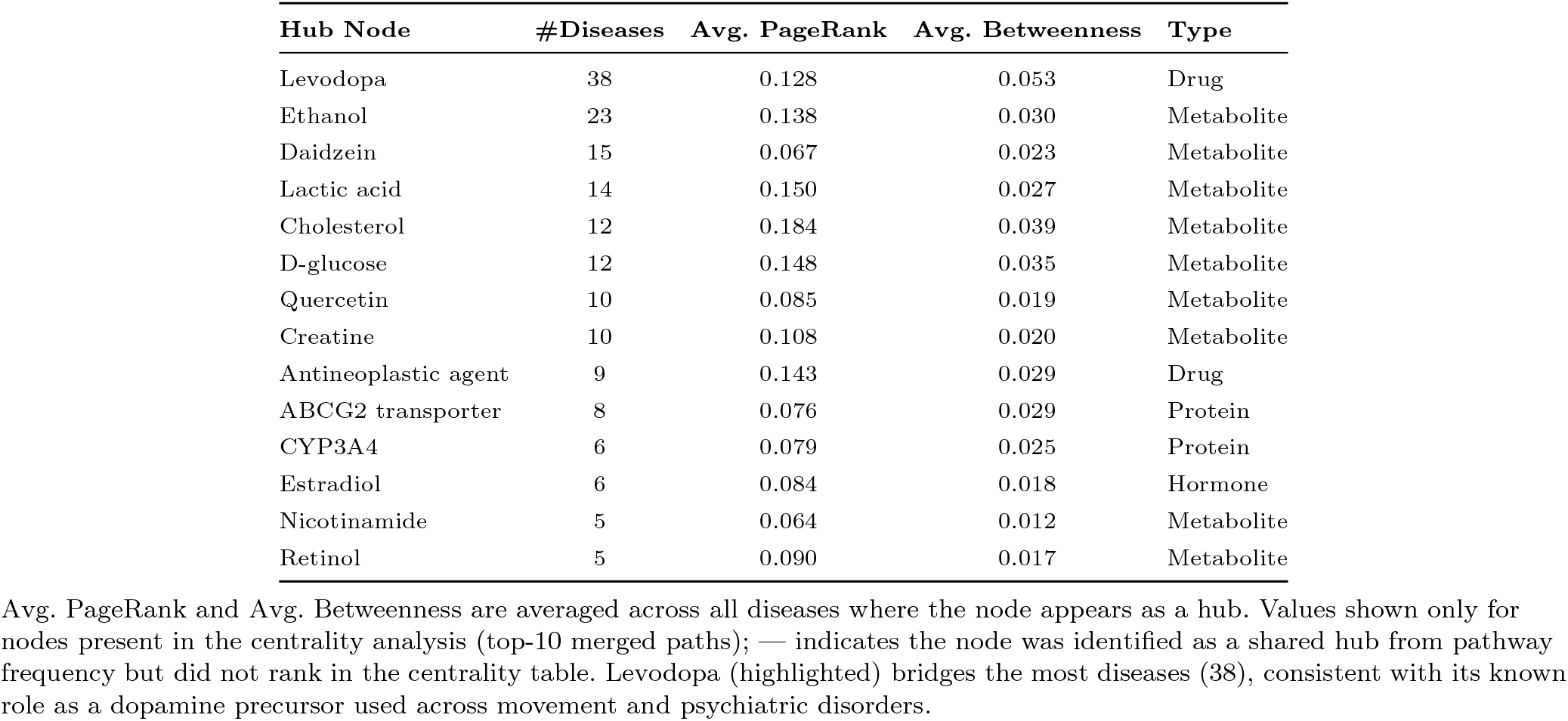
Top shared hub nodes identified across multiple diseases by graph centrality analysis. For each disease, the top-10 pathways were combined into a single graph and betweenness, and PageRank centrality were computed.

The presence of these hub structures validates our approach: key metabolites and enzymes that are known to participate in multiple biological pathways emerge as central nodes in the computationally derived pathway graphs, without any prior constraint requiring them to do so. Moreover, the identification of ABCG2 transporter and CYP3A4 as protein hubs across 8 and 6 diseases, respectively, highlights the role of drug metabolism enzymes in mediating microbe–brain interactions, a connection that warrants experimental investigation.

### GNNExplainer Stability

To assess whether the mechanistic pathways reported in this work are robust to variation, we ran GNNExplainer 5 times with different random initializations for 100 random disease– microbe pairs across 43 brain disorders. As shown in Table 5, the top-3 ranked pathways exhibited high stability: the Jaccard overlap of the top-3 paths was 0.926 *±* 0.176, and the exact same top-3 paths were ranked highest across all seeds for 83.5% of disease–microbe pairs. The top-1 pathway was consistent for 48.5% of pairs; in the remaining cases, the top-1 and top-2 paths swapped positions while the top-3 set remained stable. The near-zero path score variance (2.1 *×* 10^−8^) confirms that the GNNExplainer produces numerically consistent importance values; the occasional ranking swaps occur because adjacent paths have importance scores differing by less than 10^−4^. The high Jaccard and top-3 consistency indicate that the mechanistic pathways presented in this work are not artifacts of a single lucky initialization but represent stable outputs of the explanation method.

**Table 5.** Aggregate GNNExplainer stability metrics across 100 disease–microbe pairs (5 seeds per pair). Higher values indicate more stable explanations.

| Stability Metric | Value |
| --- | --- |
| Disease–microbe pairs analyzed | 100 |
| Seeds per pair | 5 |
| Jaccard overlap (top-3 paths) | $0.926 \pm 0.176$ |
| Top-3 path ranking consistent | 83.5% |
| Top-1 path ranking consistent | 48.5% |
| Avg. path score variance | $2.1 \times 10^{-8}$ |

### Mechanistic Pathways: Case Studies

Using the trained GNN-GBA with GNNExplainer, we identified mechanistic pathways connecting gut microbes to brain disorders. Figure 7 illustrates the top-ranked pathway for three common disorders. Below, each pathway’s mechanistic plausibility is verified from literature.

**Fig. 7.**
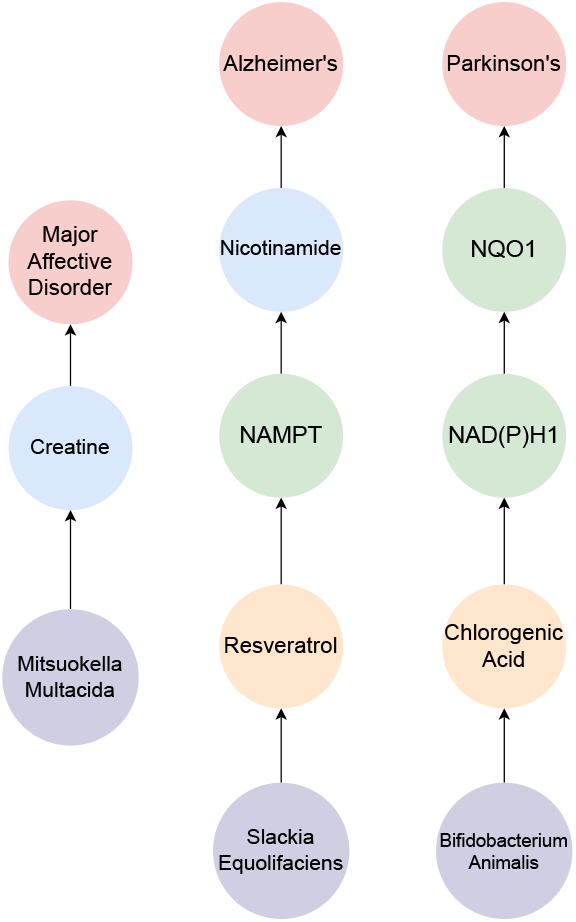
Mechanistic pathways for Parkinson’s Disease, Alzheimer’s Disease, and Major Affective Disorder.

#### Major Affective Disorder

The top pathway connecting microbes to major affective disorder (from 336 identified paths) involves Mitsuokella multacida. Mitsuokella multacida is a anaerobic bacterium found in the lower digestive tract of humans [18] that is crucial to a healthy digestive system, Mitsuokella multacida helps unlock nutrients from plant foods (via phytate degradation) and Promotes gut and metabolic health (via propionate production). [18]. While Mitsuokella multacida itself does not directly produce creatine, it is involved in arginine regulation [15], which is a precursor for creatine synthesis [27]. Creatine is primarily synthesized in the liver from arginine, glycine, and methionine [27]. Studies suggest that creatine supplementation shows promise as a potential therapy for major affective disorders [29]. Creatine plays a crucial role in brain energy metabolism, and disruptions in this process have been linked to mood disorders [29]. Research indicates that creatine may enhance the effectiveness of standard antidepressants such as selective serotonin reuptake inhibitors (SSRIs) [29]. This pathway suggests a biologically plausible mechanistic relationship between Mitsuokella multacida and major affective disorder through indirect influence on creatine metabolism, and warrants further investigation.

#### Alzheimer’s Disease

The top pathway connecting microbes to Alzheimer’s disease (from 1,455 identified paths) involves Slackia Equolifaciens. This bacterial species is found in the lower digestive tract of humans and is part of the gut microbiome, contributing to the overall microbial diversity of the digestive system [4]. Slackia Equolifaciens plays a key role in the metabolism of resveratrol, and has antioxidant and anti-inflammatory effects, and has also reported to mimic the beneficial effects of dietary restriction, extending the life span of mice and other species [4]. Moreover, resveratrol differentially regulates NAMPT [55], an enzyme, often activating it in healthy cells to craete NAD+(cellular energy) from nicotinamide and promote anti-aging pathways. Nicotinamide, a form of vitamin B3, has shown promise in preclinical studies for Alzheimer’s disease, particularly in improving cognitive function and reducing the accumulation of amyloid plaques [49]. Studies in mouse models demonstrate that nicotinamide compounds can improve cognitive function, reduce neuroinflammation, decrease DNA damage, and enhance mitochondrial function, which is often impaired in Alzheimer’s disease [3]. This pathway provides a biologically plausible mechanistic connection through which Slackia Equolifaciens may influence Alzheimer’s disease pathology, and warrants further investigation.

#### Parkinson’s Disease

The top pathway connecting microbes to Parkinson’s disease (from 1,724 identified paths) involves Bifidobacterium Animalis. This bacterium is commonly found in the gut of humans and warm-blooded animals. B. Animalis has been shown to possess enzymatic activity capable of breaking down chlorogenic acid (CGA), a common polyphenol found in plants including coffee and green tea [50]. Studies have demonstrated that B. Animalis can utilize enzymes can break down CGA into caffeic acid, allowing for the absorption of these components in the intestines [50]. Studies show that the products of the metabolism of chlorogenic acid interact with NAD(P)H dehydrogenase [quinone] 1 (NQO1) by inhibiting its activity, regulating the enzyme’s effectiveness [65, 64]. NQO1 is a crucial enzyme encoded by the NQO1 gene that plays a vital role in protecting against oxidative stress and inflammation [42]. Research suggests that dysregulation of NQO1 activity is linked to various neurological disorders, including Parkinson’s disease [38]. Impaired NQO1 function may contribute to abnormal neurotransmitter release, increased oxidative stress, and cellular damage characteristic of Parkinson’s disease [38]. This pathway provides a biologically plausible mechanistic pathway through which Bifidobacterium Animalis might influence Parkinson’s disease pathology, and warrants further investigation.

## Discussion

This work presents a comprehensive computational framework for understanding the relationship between brain disorders and the gut microbiome. By integrating diverse biomedical data into a large-scale knowledge graph (586,318 nodes, 3,573,936 edges) and applying an explainable graph neural network, we have identified specific metabolite-mediated pathways connecting gut microbes to 125 brain disorders. GNN-GBA’s decoupled architecture outperformed all nine baseline methods, and the per-relation analysis confirmed that the model maintains high ranking quality across all 103 relation types, with performance degradation limited to rare predicates with fewer than 100 test edges which is a well-understood result of class imbalance. The derived pathways align with existing literature while providing new insights into the MGBA. These pathways were not curated by hand but emerged from the model’s learned representations, scored by GNNExplainer, and shown to be stable across multiple random initializations. Beyond individual disease pathways, our analysis reveals that the MGBA operates through a small set of conserved metabolic intermediates. Flavonoids (quercetin, kaempferol, luteolin, myricetin) appear in pathways for over 114 diseases each, while bile acids (chenodeoxycholic acid, ursodeoxycholic acid) and isoflavones (genistein, daidzein) serve as shared mediators across comparably large numbers of disorders. Hub centrality analysis confirmed that nodes such as levodopa (hub in 38 diseases), cholesterol (12 diseases), and lactic acid (14 diseases) occupy structurally central positions in microbe disease pathways, with high PageRank and betweenness centrality. This suggests that therapeutic strategies targeting flavonoid bioavailability or bile acid cycling in the gut could have broadly neuroprotective effects across multiple disorders, rather than being disease-specific.

Several limitations should be noted. First, the framework is fundamentally associative: the identified pathways represent statistically plausible mechanistic hypotheses rather than experimentally confirmed results. While we validate pathway plausibility through literature consistency, the distinction between computational prediction and biological confirmation must be maintained. Second, the knowledge graph inherits biases from its source databases well-studied metabolites (e.g., quercetin, cholesterol) may appear as hubs partly because they have more recorded interactions, not solely because of their biological centrality. Third, while GNNExplainer stability is high for top-ranked paths, they degrade for lower-ranked pathways which indicates that they should be interpreted with caution.

Several extensions of this work are promising. Adding dietary components to the knowledge graph would enable GNN-GBA to identify specific foods or dietary compounds that modulate disease symptoms through the gut microbiome, supporting the development of diet recommender systems for neurological disorders. Incorporating causal inference techniques (e.g., Mendelian randomization edges, interventional data) could strengthen the mechanistic interpretability of the identified pathways beyond association. The experimental validation of the top-ranked pathways, particularly those involving the conserved flavonoid and bile acid hubs, would provide direct evidence for the computational predictions and accelerate the development of microbiome-based therapies.

The top paths for all 125 diseases are visualized in the GutBrainExplorer, an interactive dashboard that allows researchers to select diseases from a comprehensive list and visualize predicted pathways as network graphs, displaying step-by-step connections from microbes through metabolites and proteins to target diseases. Code and data are available at https://github.com/naafey-aamer/GNN-GBA.

## Author contributions statement

N.A and M.N.A conceived and conducted the experiment(s), and compiled results, and wrote the mansucript. S.V validated experiment(s) and results, and wrote the manuscript. A.D reviewed the article and performed final editing.

